# Differential regulation of GUV mechanics via actin network architectures

**DOI:** 10.1101/2022.05.01.490228

**Authors:** Nadab H. Wubshet, Bowei Wu, Shravan Veerapaneni, Allen P. Liu

**Author notes:** Corresponding author: Allen P. Liu, **Email:**. **Author Contributions:** N.H.W. and A.P.L. designed research. N.H.W., B.W. and S.V. performed research. B.W. and S.V. contributed analytic tools. N.H.W. analyzed data. N.H.W., B.W., S.V., and A.P.L. wrote the paper. **Competing Interest Statement:** Authors declare no competing interests.

## Abstract

Actin networks polymerize and depolymerize to construct highly organized structures, thereby, endowing the mechanical phenotypes found in a cell. It is generally believed that the amount of filamentous actin and actin network architecture determine cytoplasmic viscosity and elasticity of the whole cell. However, the intrinsic complexity of a cell and numerous other endogenous cellular components make it difficult to study the differential role of distinct actin networks in regulating cell mechanics. Here, we model a cell by using giant unilamellar vesicles (GUVs) encapsulating actin filaments and networks assembled by various actin crosslinker proteins. Perturbation of these cytoskeletal vesicles using AC electric fields revealed that deformability depends on lumenal viscosity and actin network architecture. While actin-free vesicles exhibited large electromechanical deformations, deformations of GUVs encapsulating actin filaments were significantly dampened. The suppression of electrodeformation of actin-GUVs can be similarly recapitulated by using aqueous PEG 8000 solutions at different concentrations to modulate viscosity. Furthermore, alpha actinin-crosslinked actin networks resulted in decreased GUV deformability in comparison to actin filament-encapsulating GUVs, and membrane-associated actin networks through the formation of dendritic actin cortex greatly dampened electrodeformation of GUVs. These results highlight the organization of actin networks regulates the mechanics of GUVs and shed insights into the origin of differential deformability of cells.

## Introduction

The cell’s ability to change shape to support cellular functions such as migration and division, and its ability to resist deformation to sustain structural integrity, depends on the cytoskeleton. Among different types of cytoskeletal polymers, actin filaments assemble into various networks aided by actin binding proteins that form large-angle-crosslinks, bundles, and branches (1,2). Although the flexural rigidity of actin filaments is as much as three orders of magnitude lower compared to that of microtubules (3), assembly of actin filaments into highly organized and dynamic networks gives rise to enhanced viscoelastic property (1,4,5). As a result, actin networks endow the mechanical phenotype of cells by differentially regulating the elasticity and cytoplasmic viscosity of cells (6). Prior research have linked the mechanical property of cells to the actin network (5,7). For example, it is reported that increased deformability of ovarian cancer cells, due to their actin organization, is directly correlated to metastatic transformation (8,9). Furthermore, retraction of epithelial cells to break cell-cell junction as a result of local actin disruption is linked to extravasation of cancer cells during metastatic invasion (10,11). It is also known that cytoplasmic viscosity of red blood cells affects their dynamics inside microvasculatures (12,13). The connection between cell mechanics and cellular processes has led to substantial interest in perturbing the cytoskeleton as a means to regulate cellular processes.

Due to the simple experimental set up, many have utilized electromechanical perturbation of cells using both direct current (DC) and alternating current (AC) electric fields. Earlier studies using electroperturbation dealt with the interaction of pulsed DC electric field and cell membranes that resulted in electropermeabilization and electroporation (14). Controlled modulation of DC electric fields resulted in the formation of enlarged pores permitting the introduction of large molecules that are otherwise not permeable through cell membrane, thus giving rise to various applications including DNA transfection, drug delivery, cancer therapy (15–17), and gene therapy (18,19). Strong AC fields, on the other hand, are known to induce cellular deformation. The semi-permeable lipid bilayer of a cell’s plasma membrane can be thought of as an electrical insulator. When an electric field is applied, ions inside a cell undergo charge separation resulting in dielectrophoresis due to a non-uniform electric field (20). Depending on the electric field strength and conductivity of the suspension environment, dielectrophoretic forces result in the deformation of cells (21). Many studies have resorted to AC electrodeformation to measure apparent stiffness of red blood cells and platelets (22,23), viscoelasticity of cancer cells (24,25), and to study the effect of actin depolymerization on the relaxation of electrodeformed cells (26).

Although prior studies have revealed the mechanical properties of cells are intimately tied to their actin networks, the differential role of actin, in the form of filaments and networks, on the deformability of cells remain incompletely understood. The intrinsic complexity of cells and numerous endogenous components make it difficult to study the differential role of actin networks as a function of actin crosslinkers (27). Giant unilamellar vesicles (GUVs) present a unique platform to model and reconstitute cellular processes in a membrane-confined environment (28). This experimental approach has been used to study the assembly of different types of actin networks (29), how actin network induces membrane remodeling (30–33), and to reveal actin binding protein competition and cooperation in actin network assembly (34,35). Others have also reconstituted a cortex-like shell in GUVs and measured their responses to mechanical compression (36).

The responses of GUVs to applied electric field have been extensively investigated (37–43). Subject to strong DC pulses, similar to cells, macropores formed in GUVs when the transmembrane potential threshold was exceeded (39). Vesicle closure after poration, curvature relaxation and other electrical properties of GUVs have been characterized for different bilayer compositions and salt concentrations used for both the external medium and the GUV lumen (37,39,44–46). Furthermore, it has been shown that when induced by a DC pulse, GUVs with an actin cortex have suppressed membrane permeability compared to cortex-free GUVs (47), presumably due to smaller and/or less macropore formation. Similar to cells, strong AC electric fields forced spherical GUVs to assume elliptical shapes with the major axis either parallel (prolate) or perpendicular (oblate) to the electric field (48). GUVs undergo these shape transformations depending on the salt concentration ratio between the GUV lumen and the solution outside of GUVs, and electric field strength and frequency (38,48,49). AC field electrodeformation transitions have been theoretically modeled under different conditions (50–52), and experimental studies based on electrodeformation have investigated bilayer properties such as membrane bending rigidity and bilayer viscosity (39,53). Although GUV membrane properties have been well studied and characterized, how the mechanics of GUV lumen affects electrodeformability remains incompletely understood.

Here, we aim to investigate the effect of encapsulated actin filaments and crosslinked actin networks on the electrodeformability of GUVs in response to AC electric fields. We encapsulated actin-free buffer solution and filamentous actin inside GUVs. Subject to an AC electric field, we observed a significant difference in deformability between the two conditions. We modulated the viscosity of GUV lumen and found that the deformability of GUVs depends on lumenal viscosity, a condition that mimics filamentous actin. Furthermore, crosslinked actin networks, in the form of lumenal network or membrane-cortex, resulted in dampened GUV deformation when compared to filamentous actin GUVs. Overall, our results reveal that the differential mechanical properties of GUVs, and by extension to cells, can be modulated by lumenal viscosity and actin network architecture.

## Results

### Actin network reconstitution in GUVs and electroperturbation device

To reconstitute various actin networks in cell-sized lipid vesicles, the modified continuous droplet interface crossing encapsulation (cDICE) method (54) (**Fig. 1A**) was used. In the presence of actin crosslinkers, actin networks formed rapidly and the modified cDICE method renders rapid encapsulation of actin networks to permit network assembly post-encapsulation. Actin filaments, components of actin cortex, and large-angle actin crosslinker (**Fig. 1B**) were encapsulated into heterogeneously sized GUVs composed of 70 mol% DOPC and 30 mol% cholesterol. 5 mol% DGS-NTA(Ni) was added when reconstituting actin cortex. Actin cortex was assembled by activating Arp2/3 complex at the inner leaflet of bilayer membrane via constitutively active His_6_-tagged VCA domain of neural Wiskott Aldrich syndrome protein (N-WASP). Crosslinked networks were formed using the large-angle actin crosslinker alpha-actinin.

**Figure 1.**
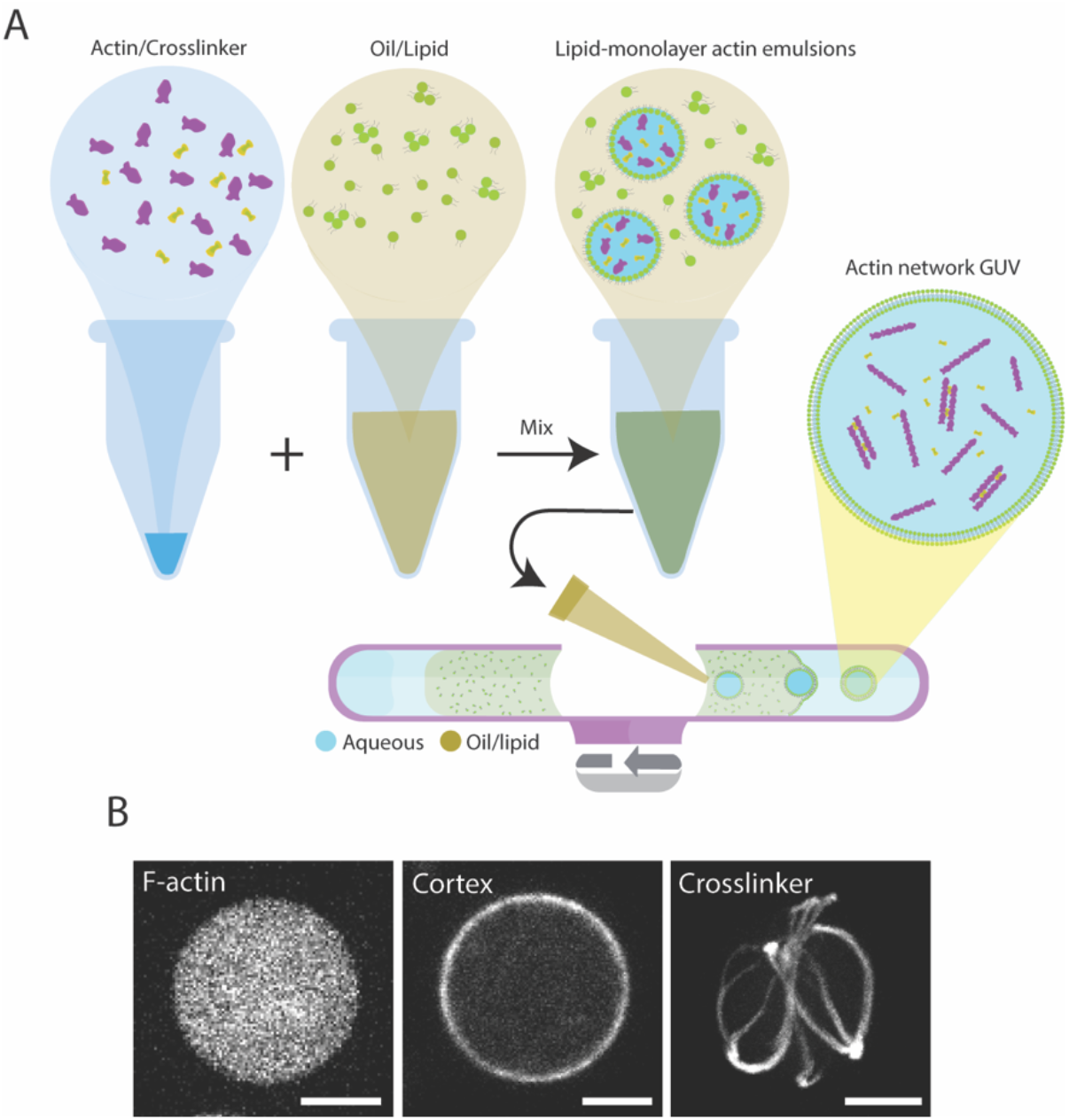
Reconstitution of different actin networks inside GUVs (A) Schematic of the modified cDICE method. Purple shapes represent actin monomers. Green shapes represent lipids. Yellow shapes, shown in actin/crosslinker solution schematic, represent an arbitrary actin crosslinker. (B) Representative images of actin network GUVs. (Left) Representative confocal image of encapsulated F-actin inside GUVs. (Middle) Arp2/3-complex assembled an actin cortex and associated to GUV lipid bilayer membrane. (Right) Aster-like actin network assembled by alpha-actinin encapsulated inside a GUV. Actin is labeled with ATTO 488 actin in all images. (Scale bars, 10 μm)

A simple electroperturbation device with two parallelly aligned and spaced electrodes was assembled on a glass slide to subject GUVs to AC electric fields at 5 kHz (**Fig. 2A**, **Fig S1A,B**). Electroperturbation experiments were performed by dispensing GUVs into the device chamber (i.e. space between electrodes), then applying a sinusoidal AC wave from a function generator. Transition of vesicles from undeformed to deformed to undeformed states following a 30 kV/m AC field at 5 kHz for a duration of 3-4 s was captured using a high-speed camera mounted on a bright field optical microscope. This setup allowed us to analyze fast real-time GUV shape transformation at a high temporal resolution.

**Figure 2.**
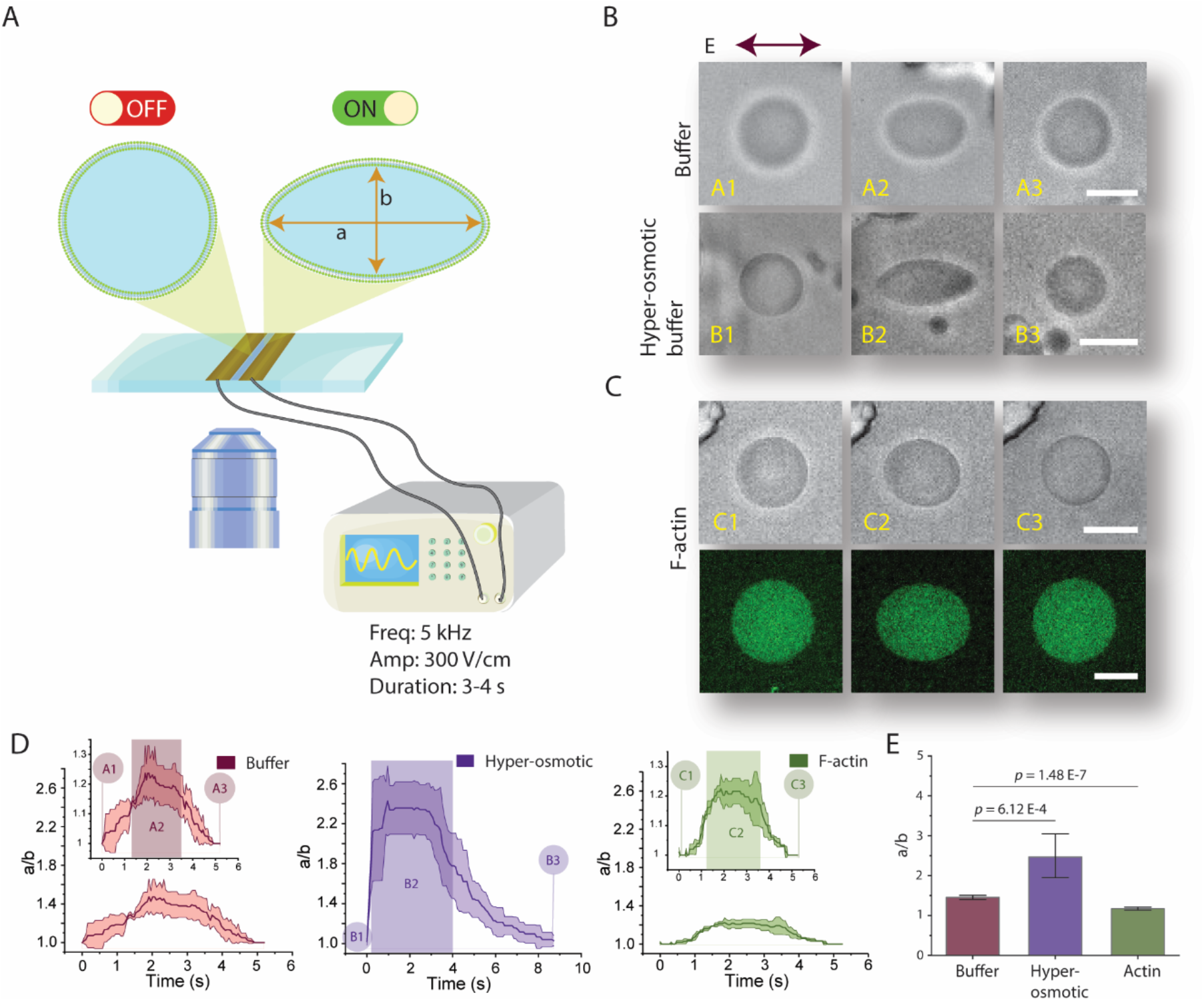
Lumenal content of GUVs alters deformation profile when GUVs are subjected to electroperturbation. (A) Schematic of the electrodeformation setup mounted on an inverted microscope. A function generator is operated at 30 kV/m at 5 kHz and a sinusoidal wave was applied for a duration of 3-4 seconds. Schematic shows electrodes adhered onto a coverslip. GUVs transform from a spherical shape to an ellipse when the electric field is applied. (B) GUV deformation is dependent on osmolarity difference between inner and outer solutions. (Top) Brightfield images of electric field-induced shape transformation of actin-polymerization-buffer GUVs. (A1) GUV at an undeformed state prior to AC field application; (A2) Steady-state deformation of GUVs during electoperturbation; (A3) actin-polymerization-buffer GUV post electrodeformation recovery. (B1, B2, B3) Electrodeformation of actin-polymerization-buffer GUV in a hyper-osmotic condition (flaccid GUV). (B2) shows exaggerated prolate deformation with pointed ends. (C) Electrodeformation of a F-actin GUV. (C2) shows visually apparent dampened deformation compared to A2 and B2. (C bottom) Representative fluorescence image of F-actin GUV labeled with ATTO 488 actin. (D) Deformation profile of GUV conditions in B and C for F-buffer, hyper-osmotic buffer, and F-actin conditions, as indicated. Labels (A1, A2, and A3…etc) correspond to GUV transformation stages during electroperturbation. Shaded rectangular box denotes approximate duration of electric field application. Shaded areas in the traces in each of the plots indicate ± SD, n = 3. (E) Comparison and statistical analysis of maximum GUV deformation of each GUV condition as indicated. Data represent mean maximum deformation and error bars denote ± SD. N_buffer_ = 11, N_hyper_ = 14, and N_actin_ = 12. (Scale bars, 10 μm)

### Actin filaments dampen electrodeformability of GUVs

First, we validated our electroperturbation setup by replicating known GUV electroperturbation responses under different ionic conditions. During AC field electroperturbation, the orientation of elliptical deformation depends on the conductivity ratio 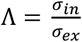, where *σ_in_* is the conductivity of inner solution and *σ_ex_* is the conductivity of outer solution (49). GUVs assume a prolate shape when Λ >1 and subjected to a low frequency field and an oblate shape when Λ <1 and subjected to a high frequency field. By tuning NaCl concentration in the inner and outer solutions and applying 30 kV/m, GUVs transformed from spherical to prorate (**Movie S1**) and spherical to oblate (**Movie S2**) shapes for Λ >1 with a low frequency field (1kHz) and Λ <1 with a high frequency field (50 kHz), respectively. Prolate deformations have major axis of ellipse parallel to the field direction, whereas oblate deformations have ellipse major axis orthogonal to the electric field.

To examine the impact of actin on GUV mechanics, we next investigated GUV deformability with and without the presence of encapsulated actin filaments (**Fig. 2**). As a control, we encapsulated actin polymerization buffer, without the presence of actin, under iso-osmotic condition. Actin polymerization buffer (hereon referred to as F-buffer) contains 1x F-buffer, 3 mM ATP, and 7.5% density gradient medium. This reaction condition contains all the reagents that are used to reconstitute filamentous actin (F-actin) and serves as the benchmark control against actin-containing GUVs. The conductivity ratio between inner and outer solution was maintained since the salt concentrations were identical for the two conditions. As expected, GUVs with F-buffer assumed prolate deformation (**Fig. 2B top**). To show that the electrodeformation is not due to GUV deflation, which can induce exaggerated deformability during electroperturbation, a control electroperturbation experiment was performed on flaccid GUVs (from hyper-osmotic condition) containing F-buffer (**Fig. 2B bottom**). This resulted in greatly increased prolate deformation of GUVs compared to their iso-osmotic counterparts. We also observed an extended delay in relaxation time for flaccid GUVs. Extended relaxation time for flaccid vesicles may be attributed to excess membrane surface area with greater membrane undulation suppressing quick recovery.

Next, we reconstituted 5.3 μM F-actin inside GUVs. Strikingly, when applying the same AC electric field to F-actin GUVs in iso-osmotic condition, deformation was significantly dampened (**Fig. 2C**, **Movie S3**). Comparing each of the above 3 conditions, the largest maximum mean deformation a/b ~ 2.42 (**Fig. 2D middle**) was attained by flaccid vesicles, followed by actin polymerization buffer GUVs, a/b ~ 1.45 (**Fig. 2D left**), and the largest deformation resistance resulted in maximum mean deformation a/b ~ 1.23 for F-actin GUVs (**Fig. 2D right**). As shown in **Figure 2E**, the average deformation from a population of F-buffer GUVs was significantly larger than that of F-actin GUVs. Considering known GUV parameters and their respective electroperturbation responses, our observation was not readily explained. In each of the above cases, there were no observed instances of electroporation to affect deformation behavior which can commonly be identified by loss of volume, loss of contrast, or by micron-sized membrane ruptures that can be observed at high magnifications. The conductivity ratio, lipid bilayer composition, and osmolarity between F-buffer GUVs and F-actin GUVs were the same. Thus, the distinct deformability behaviors can only be attributed to the material property of the GUV lumen. Previous works have shown that a F-actin solution has an increased viscosity compared to aqueous buffer solutions similar to our polymerization buffer (~1 mPa.s) and the viscosity increases further with increasing actin concentration (55). For example, Wagner *et al*. measured the viscosity of 5.3 μM F-actin solution at ~1.57 mPa.s. Thus, we hypothesized that the dampened GUV deformation was due to changes in GUV lumenal viscosity.

### Lumenal viscosity determines the electrodeformability of GUVs

In order to determine the relationship between lumenal viscosity and GUV electrodeformability, PEG 8000 solutions with concentrations ranging from 2-8% w/v were encapsulated inside GUVs. 2%, 4%, and 8% w/v PEG 8000 solutions have an estimated dynamic viscosity of 1.05 mPa.s, 3.02 mPa.s, and 6.94 mPa.s, respectively (56). Osmolarity of the outer solution was matched to the measured osmolarity of each aqueous PEG 8000 solutions in order to maintain iso-osmotic conditions. When GUVs containing 2% PEG 8000 were subjected to 30 kV/m AC field at 5 kHz for a duration of 3-4 seconds (**Fig. 3A top**, **Movie S4**), the maximum mean deformation was measured at a/b ~ 1.3 (**Fig. 3B left**). At 4% PEG 8000, the maximum mean deformation reduced to a/b ~ 1.14 (**Fig. 3A middle**, **Fig. 3B right**, **Movie S5**). In GUVs with 8% PEG 8000, no measurable GUV deformation was observed (**Fig. 3A bottom**, **Movie S6**). Each GUV population at variable encapsulated PEG concentrations was significantly different from each other (**Fig. 3C**), and clearly demonstrated dampening of GUV deformation with increasing lumenal viscosity.

**Figure 3.**
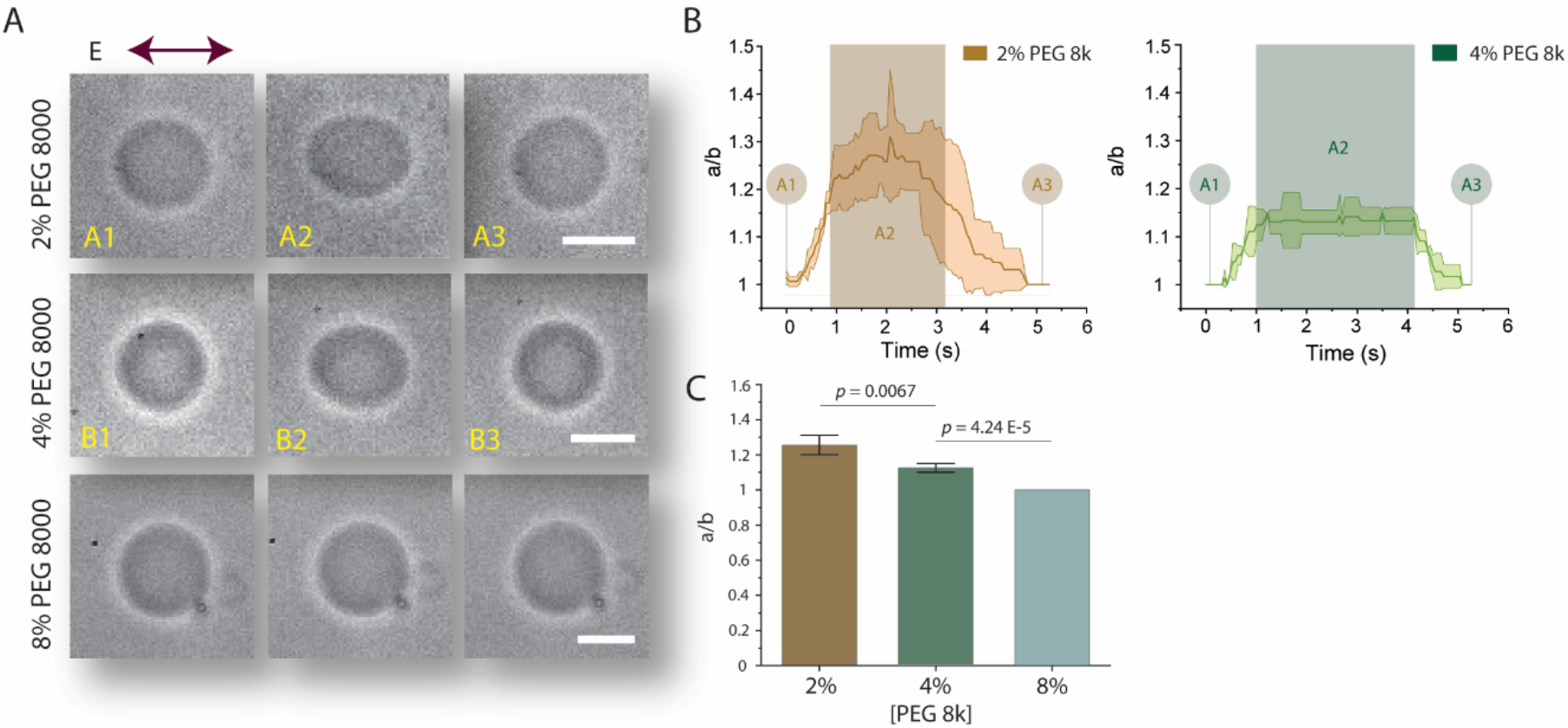
Relationship between viscosity contrast and electrodeformability of GUVs. Different PEG 8000 concentrations were encapsulated inside GUVs to vary viscosity contrast between GUV lumen and outer solution. (A) Brightfield images showing PEG 8000 GUV shape transitions from undeformed to elliptically deformed to spherical recovery. (Top) Electroperturbation of 2% w/v PEG 8000 GUV. (A1) A PEG 8000 GUV at an undeformed state prior to AC field application. (A2) Steady-state deformation of GUVs during electoperturbation. (A3) GUV post electrodeformation recovery to assume spherical shape. (B1-B3) 4% w/v PEG 8000 GUV electrodeformation. (Bottom) Electroperturbation of 8% w/v PEG 8000 GUV. (B) Deformation profile of 2% and 4% w/v PEG 800 GUVs, n = 3. Shaded rectangular box denotes approximate duration of electric field application. Shaded areas in the traces in indicate ± SD. (C) Comparison and statistical analysis of maximum GUV deformation of each GUV conditions indicated. Data represent mean maximum deformation and error bars denote ± SD. N_2%=_ 13, N_4%_ = 11, and N_8%_ = 12. (Scale bars, 10 μm)

Our results are consistent with our initial hypothesis that dampened deformation in Factin GUVs is possibly due to a change in viscosity. The degree of deformability of 5.3 μM F-actin GUVs falls between deformability of GUVs encapsulating 2% PEG and 4% PEG solutions, and directly corresponds to the measured viscosity of F-actin solution at this concentration (1.5 mPa.s) that is in between the viscosity of 2 and 4% PEG solutions. Although these observations may be intuitive and in alignment with our initial hypothesis, to our knowledge, there are no prior studies that exploited cell-mimicking confinements like GUVs to investigate the effect of lumenal viscosity on their electrodeformability. Thus, here we illustrate a mechanism for cells to maintain structural integrity by only modifying viscosity without introducing structural complexity of actin networks in cell mimetics.

### *In silico* investigation on the role viscosity contrast on GUV electrodeformability

We developed a computational method to further investigate the role of viscosity contrast (detailed in SI). Numerical experiments are set up by placing the GUV in an AC field *E*_∞_(*t*) with magnitude *E*_0_, such that

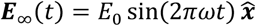

where ω is the AC field frequency. Using the GUV radius *a* as the characteristic length scale and the membrane charging time *t_m_* = *aC_m_/σ_ex_* as the characteristic time scale, we define the following dimensionless parameters:

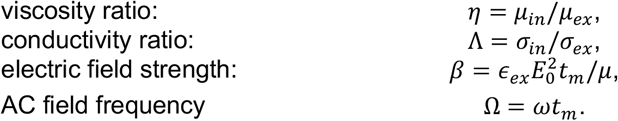

An AC field of frequency Ω = 0.5 and strength *β* = 10 is applied at *t* = 0 and we measure the aspect ratio a/b of the GUV over time. We observe that, for a fixed conductivity ratio Λ, the prolate deformation of the GUV is delayed as the viscosity contrast *η* is increased (**Fig. 4A**). Additional experimental results, shown in **Figure S2,** indicate that fixing the conductivity ratio *Λ* of 2% (**Fig. S2A**) and 4% PEG 8000 (**Fig. S2B**) concentrations to 0.9, by addition of 7.5 mM NaCl to 4% PEG 8000 inner solution, preserved deformation dampening as a result of increasing *η* (**Fig. S2C,D,E**). For a fixed viscosity contrast, the prolate deformation happens only when the conductivity ratio Λ is large enough (**Fig. 4B**). Consequently, dampening of the prolate deformation is observed as a combined effect of increasing *η* and decreasing Λ (**Fig. 4C**), which is consistent with the experimental results using PEG 8000 solutions with increasing concentrations (**Fig. 3**).

**Figure 4.**
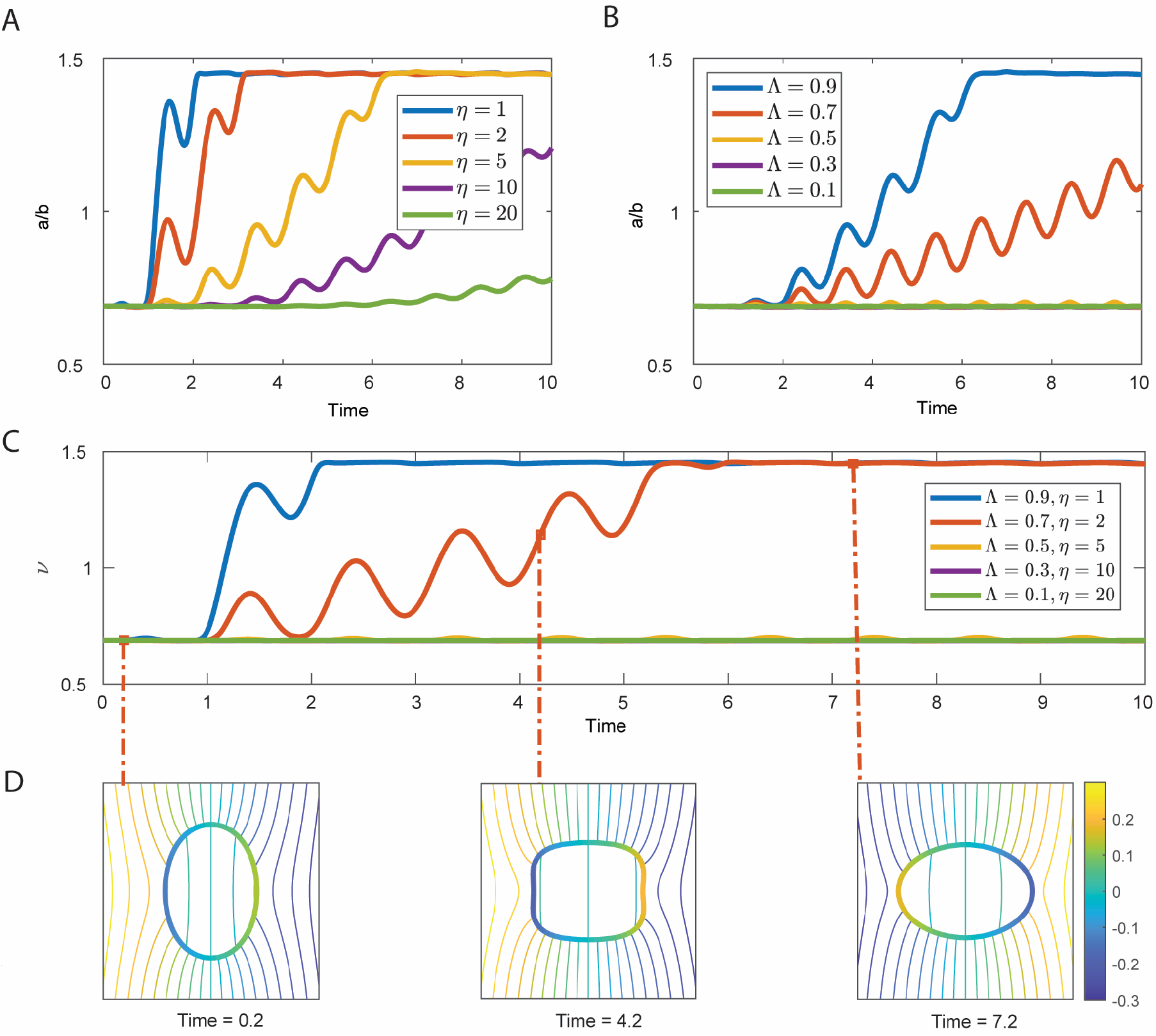
Prolate deformation of a GUV suspended in an AC field (*β* = 10, Ω = 0.5) obtained via numerical simulations. (A) Electrodeformation for various viscosity contrasts *η* while the conductivity ratio is fixed (Λ = 0.9). We observe that the higher GUV luminal viscosity, the longer the time it takes to complete the prolate deformation. (B) Conductivity ratio is varied while the viscosity contrast is fixed to *η* = 5. Prolate deformation takes longer as Λ is reduced and halts altogether below a threshold Λ. (C) Decreasing Λ and increasing *η* simultaneously results in a compounding effect on the prolate deformation, which is highlighted in this experiment. (D) Electric potential contour plots around the vesicle of Λ = 0.7, *η* = 2 at times *t* = 0.2 (flaccid GUV) *t* = 4.2 (transitionary phase) and *t* = 7.2 (prolate).

One advantage of the numerical approach presented in this section is its ability to collect a variety of quantities of interest that will be useful for further investigations. For example, we have shown in Figure 4D the contour plots of the electric potential around the vesicle that corresponds to different stages of the deformation, where it is clear that when the vesicle is being stretched during the transitioning stage, the electric field strength is almost uniformly zero inside the GUV.

### Structurally distinct actin networks differentially regulate GUV mechanics

Mechanical features and responses of actin networks are governed by actin binding proteins and particularly actin crosslinkers. These crosslinkers not only assemble phenotypically distinct networks but also spatially organize actin networks allowing the cell to have variable mechanics across the cell volume. How might structurally distinct actin networks in a cell-mimicking confinement determine mechanical behavior? Here, we examined GUVs with Arp2/3-branched dendritic actin cortex (actin-cortex GUVs) or networks made with a large-angle actin crosslinker alpha-actinin (alpha-actinin-crosslink GUVs). Membrane-bound dendritic actin cortex was achieved by activation of Arp2/3 complex using membrane-associated nucleation promotion factor (His_6_-tagged VCA) on DGS NTA(Ni)-containing membrane. Encapsulating actin with His_6_-tagged VCA-activated Arp2/3 complex generated uniform actin-cortex GUVs with a high efficiency (**Fig. 5A top**), whereas alpha-actinin addition led to a range of actin network morphologies, including rings, asters and random networks (**Fig. 5A bottom**). Although reconstitution of various actin networks is well established to study crosslinkers and network phenotypes (34,35,57,58), little is known about how these actin crosslinkers differentially regulate GUV deformability.

**Figure 5.**
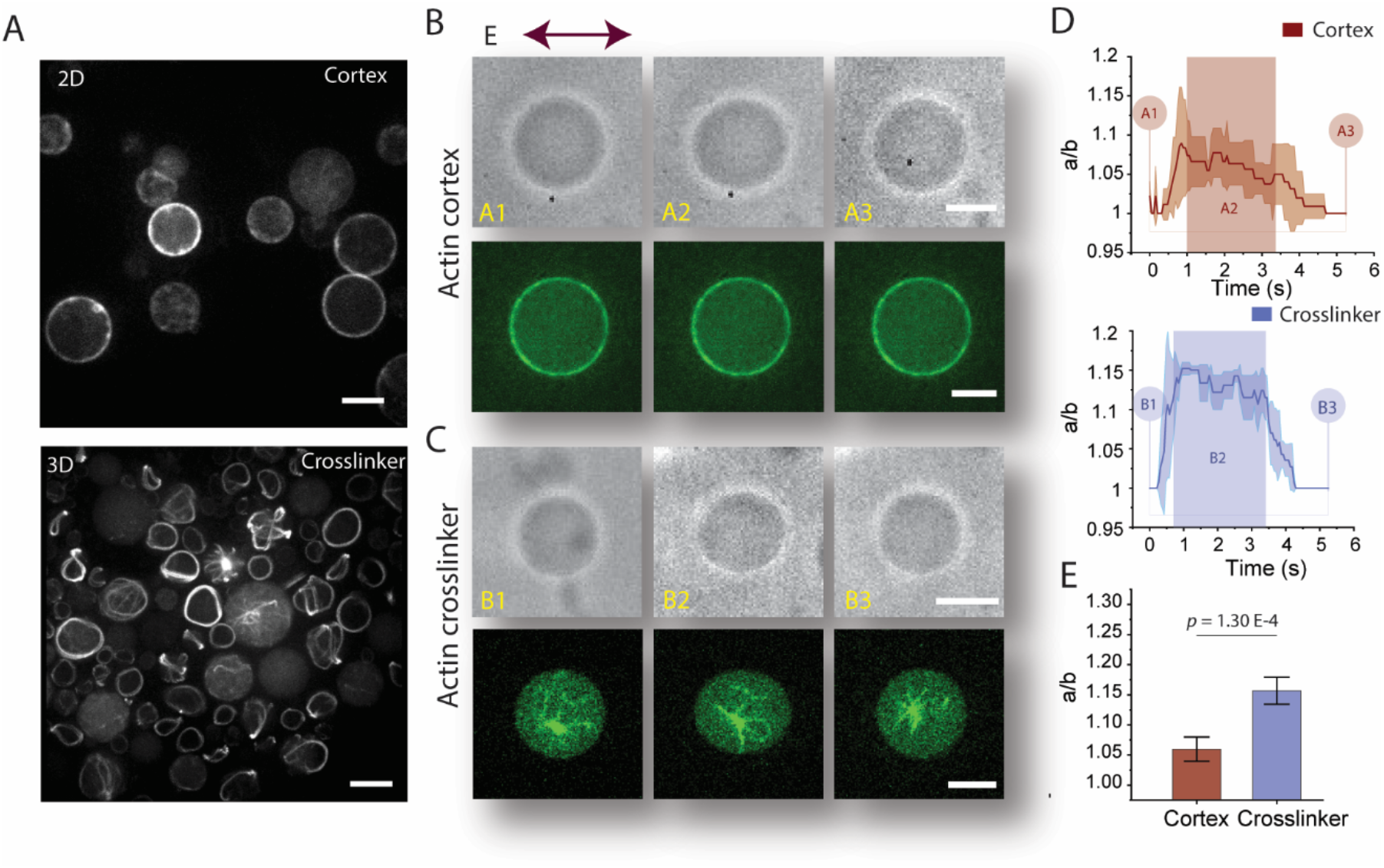
Actin networks reduce deformation induced by AC field electroperturbation. (A) High efficiency reconstitution of actin-cortex GUVs and alpha-actinin-crosslink GUVs using the modified cDICE method. (Top) Representative confocal image of Arp2/3 complex-assembled dendritic-actin-cortex GUVs. GUVs have a uniform actin cortex shell associated to the membrane via His_6_-tag-nickel interaction. (Bottom) Representative confocal image of alpha-actinin-crosslink GUVs. Various actin network phenotypes commonly seen with actin networks with large-angle crosslinkers were observed. Both images show ATTO 488 actin to label actin networks. (B) Electroperturbation of actin-cortex GUV. (Top) Brightfield images showing shape transitions pre (A1), during (A2) and post (A3) application of AC electric field. (Bottom) Confocal images of ATTO 488 actin showing actin cortex GUVs corresponding to A1, A2, and A3. (C) Electroperturbation of alpha-actinin-crosslink GUV. (Top) Brightfield images showing shape transitions pre (B1), during (B2) and post (B3) application of AC electric field. (Bottom) Confocal images of ATTO 488 actin showing alpha-actinin-crosslink GUVs at different stages of electroperturbation. (D) Deformation profile of actin-cortex GUVs (Top) and alpha-actinin-crosslink GUVs (Bottom) GUVs. n = 3. Shaded rectangular box denotes approximate duration of electric field application. Shaded areas in the traces in indicate ± SD. (E) Statistical analysis of electrodeformed actin-cortex and alpha-actinin-crosslink GUVs. Data represent mean maximum deformation and error bars denote ± SD. N_cortex_ = 12 and N_crosslink_ = 11. (Scale bars, 10 μm)

We followed the same electroperturbation procedure employed in previous experiments and subjected actin cortex GUVs to an AC field. Electrodeformability of actin-cortex GUVs was greatly dampened and hardly visible to the naked eye (**Fig. 5B**). Under the same condition, GUVs with alpha-actinin-crosslinked networks were more deformed compared to actin-cortex GUVs (**Fig. 5C**). Electrodeformation was dampened to the largest extent by actin-cortex GUVs with a max mean deformation a/b ~ 1.07 (**Fig. 5D top**), and alpha-actinin-crosslink GUVs had a max mean deformation of a/b ~ 1.15 (**Fig. 5D bottom**). Looking closely at the deformation profile of actin-cortex GUVs, strangely, as shown in Figure 5D top, deformation was not sustained over the duration of AC field application, but rather GUVs started to recover immediately after reaching max deformation. Similar profile was not observed for alpha-actinin-crosslinked GUVs as they maintained deformation throughout the duration of applied AC field.

Compared to the F-actin GUVs, both actin-cortex and alpha-actinin-crosslink GUVs had reduced deformability, with extreme dampening in actin-cortex GUVs (**Fig. 5E**). These results demonstrated differential mechanical properties of various isolated actin networks in a cell-like confinement. The mechanism by which actin networks achieve such mechanical variation is still open to investigation. Although prior studies reconstituting various bulk actin networks *in vitro* have shown that crosslinking actin filaments results in enhanced viscosity (55), we cannot, however, due to the complexity of actin network organization, directly and fully attribute these differences in deformability to lumenal viscosity induced by actin network assembly.

## Discussion

In this work, we examined how the material property of lumenal contents determines electrodeformability of cell-mimicking GUVs subjected to an AC electric field. This mechanism is distinct from conditions that are known to impact the degree of electrodeformation which include conductivity contrast, osmotic contrast, lipid bilayer composition, lipid bilayer viscosity, and electric field intensity (40,49,50,53). In contrast, our results show that deformability of GUVs encapsulating actin filaments is suppressed compared to actin-free GUVs. We demonstrated the deformation dampening of F-actin GUVs is due to increased viscosity of GUV lumen. Motivated by differential mechanics found in a cell, we further examined how actin cortex and crosslinked actin networks govern GUV electrodeformation. Evident from dampened deformation, our results illustrate that actin network GUVs have enhanced mechanical resistance, and that there is also GUV deformability difference between different architectures.

As revealed by our findings, actin-cortex GUVs have greater deformation resistance compared to GUVs with alpha-actinin-crosslinked networks. Prior findings by Wagner *et al*. show that actin crosslinkers increase viscosity of actin in bulk solutions (55). However, the structure and spatial scale of actin networks formed in bulk solutions are diametrically different from those assembled in a cell-like confinement. Thus, it would be premature to attribute our finding that actin crosslinkers differentially regulate GUV electrodeformability to simply viscosity difference. During electrodeformation, GUVs undergo two distinct deformation regimes namely entropic and elastic regimes (42). The extent of the deformation in the entropic regime is dependent on the degree of thermal undulations in the bilayer which varies depending on the lipid composition and osmotic contrast, whereas the elastic regime is dictated by field intensity and bilayer stretchability at the molecular level. These deformation regimes may potentially be altered as a result of the material property of the lumen and its interaction with the lipid bilayer membrane. Thus, it is important to consider mechanisms of how different actin networks may affect these deformation regimes beyond changes in lumenal viscosity.

It is well established that the actin cortex regulates membrane rigidity (59,60). When thin actin-cortex shells were reconstituted in GUVs and subjected to hydrodynamic tube pulling, it was shown that the membrane tube length was reduced for thin actin shell GUVs (61). Considering this prior finding, it is possible that the mechanism of electrodeformability suppression by actin cortex is due to changes in membrane rigidity that restricts membrane undulation in the entropic regime of deformation, and also restricting lipid mobility, consequently reducing bilayer stretching, in the elastic regime of electrodeformation. This and the overall change in lumenal viscosity due to presence of F-actin and crosslinker may collectively contribute to the increased deformability resistance of actin-cortex GUVs. For alpha-actinin-crosslink GUVs, a different mechanism may be plausible to account for their suppressed electrodeformability. When F-actin GUVs are subject to an electric field, due to the scale of individual filaments with respect to GUV size and field pressure, actin filaments are unable to individually resist deformation, akin to sand grains in quicksand, and unable to undergo individual strain. However alpha-actinin assembles complex actin scaffolds that are capable of reinforcing the GUV, like a truss system, to resist field forces. Further investigation could possibly shed more light on the relationship between crosslink/bundle rigidity and electrodeformability.

For the numerical simulation of the electrodeformation of GUVs, the leaky-dielectric model is used, which characterizes some key physical and mechanical properties of GUVs, including conductivity contrast, membrane rigidity, and lumenal viscosity. Our numerical simulations provide additional supporting evidence, independent of experiments, that GUVs of increased lumenal viscosity experience greater deformation resistance. An important advantage of numerical simulation is its ability to collect various quantities of interest at ease, such as electric potential and velocity fields, offering more detailed characterizations of GUV electrodeformation. However, there are limitations to the current mathematical model. Firstly, due to its simplifying assumptions on the membrane structure, this model is incapable of capturing phenomena such as electropermeabilization and electroporation that occur under a strong electric field. Thus, restricting the membrane to an inextensible and intact boundary. Secondly, the current model does not account for the cytoskeleton structures in GUVs, which will be important for further investigating the effect of different actin networks as integral structural components on the electrodeformability of GUVs. More sophisticated mathematical models need to be developed.

There are many fascinating mechanobiological inquiries that can be pursued using cytoskeletal GUVs. The cell is a very dynamic and structurally and functionally complex system with many proteins involved in a single function. The GUV furnishes a cell-like confinement system that is suitable for systematic construction of complex cellular functions module by module. Using our findings as a steppingstone, we anticipate future interest in examining the role of various other types of actin networks, and co-assembled networks of actin, intermediate filaments, and microtubules, in determining mechanophenotypes. Such efforts will help uncover deep insights into cell mechanics from the bottom up.

## Materials and Methods

### Reagents

Purified actin was purchased (Cytoskeleton, Inc.). ATTO 488 actin was purchased from Hypermol (Germany). Actin crosslinker alpha-actinin from rabbit skeletal muscle and Arp2/3 complex from bovine brain were purchased form Cytoskeleton, Inc. Hexa-histidine-VCA (His_6_-tag VCA) was purified as described previously (35). General actin buffer (G-buffer) was prepared at 10x concentration and consists of 50 mM Tris-HCL pH 8.0, and 2 mM CaCl_2_. Actin was diluted from a stock concentration of 10 mg/ml to a working concentration using G-buffer + 0.2 mM ATP and 0.5 mM DTT. Actin polymerization buffer (F-buffer) was prepared at 10x concentration and is composed of 500 mM KCl, 20 mM MgCl_2_, and 10 mM ATP. 1,2-dioleoyl-sn-glycero-3-phosphocholine (DOPC), 1,2-dioleoyl-sn-glycero-3-[(N-(5-amino-1-carboxypentyl)iminodiacetic acid)succinyl] (nickel salt) (DGS-NTA(Ni)), and cholesterol were purchased from Avanti Polar Lipids. PEG 8000 was purchased form Fisher Scientific. Density gradient medium (Optiprep) and other chemicals were purchased from Sigma Aldrich.

## Acknowledgments

We thank Yashar Bashirzadeh for helpful discussion. The work is supported by the National Science Foundation (CBET-1844132). N.H.W. was supported by NIH’s Microfluidics in the Biomedical Sciences Training Program (NIH NIBIB T32 EB005582). S.V. acknowledges support from National Science Foundation (DMS-2012424). A.P.L. acknowledges support from National Institutes of Health (R01 EB030031-01) and National Science Foundation (EF1935265 and MCB220136).

