## Supplemental Information for "Differential regulation of GUV mechanics via actin network architectures"


**This PDF file includes:**

Supplementary text
Figures S1 to S2
Tables S1 to S2
Legends for Movies S1 to S6
SI References

**Other supplementary materials for this manuscript include the following:**

Movies S1 to S6

**Supplementary Information Text**

**Materials**

**Electrodeformation setup**

A simple homemade electroperturbation chamber was assembled to conduct GUV electrodeformation experiments. The setup is comprised of adhesive electrode tape and 3 x 1 in coverslip. Two copper tapes were used as electrodes and were attached to one face of the coverslip in a parallel manner with gaps ranging from 200-300 µm. Sinusoidal AC electric field was applied using an Agilent 33120A (Keysight Technologies, USA) function generator between electrodes adhered to coverslip. We used a fixed length of electrode tapes, and using voltmeter (Fluke, USA), we measured potential of 6.7 V when applying 10 V peak to peak sinusoidal AC. Lower than RMS voltage measured at the electrodes end may be attributed to resistance from adhesive and tape length. To apply identical AC field between slightly varying chambers between different devices, applied voltage was adjusted accordingly to the exact measured gap between the two electrodes such that 30 kV/m is the applied field strength between the electrodes. During electroperturbation experiments, GUVs were dispensed between the electrodes. The height of the copper tape, which is ~100 µm, was higher than nearly all sizes of generated and analyzed GUVs, therefore yields an uniform electric field across the length of the chamber. For all experiments, duration of AC field was kept within 3-4 seconds.

**GUV generation**

Encapsulation of aqueous material inside GUVs was achieved using the modified cDICE method (1). As described previously (2), a 3D printed cDICE chamber is mounted onto a table top stirring motor and rotated at 1200 rpm. First, 770 µL outer aqueous glucose solution, of varying concentrations depending on osmotic condition, is dispensed into the chamber. For iso-osmotic conditions, concentration of glucose is tuned such that its osmolarity matches the measured osmolarity of the inner solution, whereas for hyperosmotic conditions (flaccid GUVs), the outer glucose solution is 400 mOsm higher than inner solution. Next an adequate amount of oli/lipid mixture is dispensed into the chamber. The lipid composition used in all conditions, except for reconstituting actin cortex, is 70 mol% DOPC with the addition of 30 mol% cholesterol. During reconstitution of the actin cortex, 5 mol% DGS NTA(Ni) was added to the lipid composition while lowering cholesterol to 25 mol%. Oil is composed of 80% silicon oil and 20% mineral oil. When oil and lipid solutions are mixed, a two-phase dispersion emerges due to the emulsification of mineral oil containing lipid aggregates. Upon the addition of the lipid/oil mix to the chamber, it forms an interface saturated by lipid aggregates. Separately, 770 µL of oil/lipid mix is dispensed in to an epitube containing 20 µL of prepared inner solution (encapsulant) and pipetted up and down until the solution becomes cloudy indicating formation of lipid monolayer saturated encapsulant emulsions. Finally, the solution is transferred to the cDICE chamber. Due to centrifugal forces generated by the rotating chamber, encapsulant emulsions are shuttled through the oil/lipid mix into the outer solution. When emulsions cross the lipid saturated interface, a second layer of lipid zips the emulsions and forms GUVs suspended in the outer aqueous solution.

**Inner solution preparation**

Various inner solution conditions were reconstituted to conduct GUV electroperturbation experiments. Each condition contains 7.5% density gradient medium to facilitate GUV sedimentation. In viscosity contrast experiments, PEG 8000 was dissolved in DI water at specified concentration (2%, 4%, and 8% w/v). To reconstitute actin-polymerization-buffer GUVs, inner solution contained 1x F buffer and 3 mM ATP. For reconstitution of F-actin GUVs all components in the inner solution of actin-polymerization-buffer GUVs are preserved with the addition of 5.3 µM actin and 0.53 µM ATTO 488 actin. To reconstitute alpha-actinin-crosslink GUVs, all ingredients used to reconstitute F-actin were mixed and incubated for 15 minutes in ice. Then, 1.77 µM of alpha-actinin was added to the solution. Addition of alpha-actinin, or any actin crosslinker, should be immediately followed by the last step of the cDICE GUV generation method, which is making lipid monolayer stabilized inner solution emulsions by mixing actin solution with lipid/oil mix followed by dispensing into the cDICE chamber. For reconstitution of actin cortex, the lipid composition is slightly altered by the addition of 5 mol% DGS NTA(Ni). Similar to alpha-actinin-crosslinked actin network reconstitution, F-actin components were incubated in ice for 15 minutes. Then actin nucleation promotion factor, 0.5 μM $His_6$-tagged VCA, is added followed by addition of 0.5 μM of Arp2/3 complex. When confined by the lipid bilayer compartment, $His_6$-tagged VCA binds to nickel domain of DGS NTA(Ni) and activates Arp2/3 to form dendritic actin networks restricted at the lipid bilayer membrane.

#### Imaging

In our experiments, we used two different imaging setups: bright-field imaging equipped by a high-speed camera and fluorescence imaging using a confocal microscope. To acquire dense data points yielding contentious deformation profile of GUVs when subject to AC electric field, Olympus CKX41 (Olympus, USA) inverted microscope equipped with Phantom Miro ex1 (Phantom High Speed, USA) high-speed camera was used. Images were taken at a rate of 1200 pps using a 40x/0.55 NA objective lens and acquired using phantom camera control (PCC 1.2) software. To observe actin dynamics in response to electric field, we used an Olympus IX-81 inverted microscope equipped with a spinning disk confocal (Yokogawa CSU-X1), OBIS LS/LX lasers (Coherent, USA) and an iXON3 EMCCD camera (Andor Technology, USA). Each component was controlled by using MetaMorph (Molecular Devices, USA). Images were acquired using an oil immersion 40x/1.3 NA objective. While GUV samples were inside the electroperturbation chamber and subject to an AC electric field, ATTO 488 actin was excited using 488 nm laser at an exposure time of 170 ms, and time-lapse images were taken every 200 ms. Maximum deformation measured from confocal images were, along with bright-field images, used for statistical analysis of each GUV condition.

### Methods

#### Image processing

Images acquired from high-speed camera equipped bright-field microscopy are saved as a Phantom Cine File (.cine). Since images acquired from PCC 3.6 are taken at a high rate, number of images are reduced by decimating frames as needed. Images are further processed using ImageJ. To import .cine files, Cine File Importer (CINE_File.jar) add-in is included in the ImageJ plugins list. Reduced .cine images are then imported to analyze deformation profile of GUVs frame by frame. To do this, image brightness and contrast were adjusted such that there is a distinct and clear boundary between GUV boundary and outer solution. Depending on the GUV condition, contrast of raw GUV images are not consistent from one image to another, thus *Brightness/Contrast* adjustment must be tweaked on a case by case basis. Then, images are transformed to binary pixel classes using the *threshold* menu in ImageJ. After thresholding images, clear boundaries are outlined and an ellipse is fitted onto GUVs using the *Fit ellipse* command. Major and minor axis of fitted ellipses were selected as measurement parameters and reported.

#### Numerical method

The parameters used in this numerical analysis are summarized in **Table S1**. Consider a GUV comprised of charge-free bilipid membrane with its interior and exterior filled with a fluid of viscosities $\mu_{in}$ and $\mu_{ex}$ respectively. To model the electrohydrodynamics, we will employ the leaky dielectric model (3), which combines the Ohm's law for electric current conservation and the Stokes equations for fluid motion. The fluid velocity u satisfies

$$-\mu\nabla p + \Delta \boldsymbol{u} = 0, \quad \nabla \cdot \mathbf{u} = 0,$$

in the interior and exterior of the vesicles subject to a far-field condition and a no-slip boundary condition at the GUV boundary γ. In addition, at γ, the membrane elastic forces balance the electric and hydrodynamic forces, that is, $\mathbf{f}_{mem} = \mathbf{f}_{el} + \mathbf{f}_{hd}$. The membrane elastic forces are obtained by taking the gradient of the Helfrich energy, $E_m = \frac{1}{2}\left(\int_\gamma \kappa_b \kappa^2 d\gamma\right)$, that is used for modeling the membrane energy. Here, $\kappa_b$ is the bending modulus and κ is the planar membrane curvature. The local inextensibility of the membrane is enforced by letting the surface divergence of the interfacial velocity vanish, that is,

$$\nabla\gamma \cdot \dot{\boldsymbol{x}} = 0,$$

where **x** is assumed to be the position of the interface. This constraint will be enforced via augmented Lagrangian approach. Thereby, it gives rise to an additional interfacial force due to tension λ, the Lagrange multiplier. The combined expression is given by,

$$\boldsymbol{f}_{el} = -\kappa_b\left(\kappa_{ss} + \frac{\kappa^3}{2}\right)\boldsymbol{n} + (\lambda\boldsymbol{x}_s)_s,$$

where **n** is the outward normal to vesicle interface. The remaining component we require to close the system of equations for vesicle EHD is the electric force $\mathbf{f}_{el}$ acting on the fluid. It is given by the jump in the normal component of the Maxwell stress tensor:

$$\boldsymbol{f}_{el} = \left[\left[\boldsymbol{n}\cdot\left(\epsilon\boldsymbol{E}\otimes\boldsymbol{E} - \frac{1}{2}\epsilon\left|\left|\boldsymbol{E}\right|\right|^2\boldsymbol{I}\right)\right]\right],$$

where ε is the permittivity, **E** is the electric field and [·] is the difference between interior and exterior fields. The ambient electric field is conservative and can be computed from the electric potential, **E** = −∇φ, by solving the Laplace equation, −∆φ = 0, in the interior and exterior of the vesicle interface. The boundary conditions at the fluid-membrane interface are obtained by charge and current conservation across the membrane (4). The charge accumulation is governed by: (i) Charge convection by the fluid motion along the surface, (ii) Membrane conductance, with strength $G_m$, arising from the presence of pores, pumps and ion channels. (iii) Membrane capacitance $C_m$. Together, the interfacial conditions can be written as

$$\left[\left[\sigma E_n + \epsilon\dot{E}_n\right]\right] = 0$$

$$C_m\dot{V}_m + G_mV_m = \sigma_{ex}E_{n,ex} + \epsilon_{ex}\dot{E}_{n,ex}$$

where σ is the fluid conductivity, $E_n$ is the normal electric field at the membrane interface and $V_m$ = [[φ]] is the potential difference across the membrane. The values used for this numerical analysis are summarized in **Table S2**.

In summary, given the initial shape of a GUV, we need to solve for the electric potential and the fluid velocity at the interface, advance the interface position via the kinematic condition, and update the membrane electric variables using (5). We employ the boundary integral formulation developed in (6) for solving the Stokes equations and that of (5,7) for the electric potential problem, with appropriate modifications to account for the imposed AC electric field (as opposed to DC field considered in those works).

**Statistical analysis**

Using Origin software**,** one-way ANOVA tests were used to determine the statistical significance of major axis to minor axis ratios across different conditions. Furthermore, *p* values were determined using two-tailed Student's *t*-test. $p < 0.05$ is considered statistically significant. Number of vesicles measured range from 10 to 15 with at least three different devices used for a given condition.

### Dimensionless parameters used for numerical simulation

**Outer solution property**
$\varepsilon_{ex}$ of 200mM glucose = 79.4 (8) absolute $\varepsilon_{ex}$ = 7.03 x $10^{-10}$
$\sigma_{ex}$ of 200 mM glucose = 0.179 mS/m (8)
$\mu_{ex}$ of 200 mM glucose = 1mPa.s (9)

**Membrane property**
$C_m$ = 1uF/cm^2 (10)
A ~10um
$G_m$ = 0, assuming intact lipids (10)
κ = $10^{-19}$ J

**Applied Electric field**
$E_o$ = 30 kV/m
ω = 5 kHz

**Inner solution property (PEG8000 2%, 4%, 8%)**
$\varepsilon_{in}$ PEG8000 = 80.2, absolute $\varepsilon_{ex}$ = 7.1 x $10^{-10}$
$\mu_{in}$ of 2% PEG = 1.05 mPa.s
$\mu_{in}$ of 4% PEG = 3.02 mPa.s
$\mu_{in}$ of 8% PEG = 6.94 mPa.s (11)

$\sigma_{in}$ of 2% PEG = 16.7 dS/m
$\sigma_{in}$ of 4% PEG = 14.1 dS/m
$\sigma_{in}$ of 8% PEG = 11.7 dS/m (12)

Electrical conductivity values of aqueous PEG 8000 solutions were acquired from Burnett et. al. (12). In this article, electrical conductivity of PEG 8000 was measured for various PEG 8000 concentrations in Hoagland solution. Within the range of 0-10% w/v PEG8000 concentration, electrical conductivity was measured to have a linear correlation with PEG8000 concentration. To calculate the conductivity of PEG 8000 dissolved in water, we linearly interpolated for unknown values of x% w/v PEG8000 electrical conductivity in water using electrical conductivity of water and Hoagland solution as the independent variables.

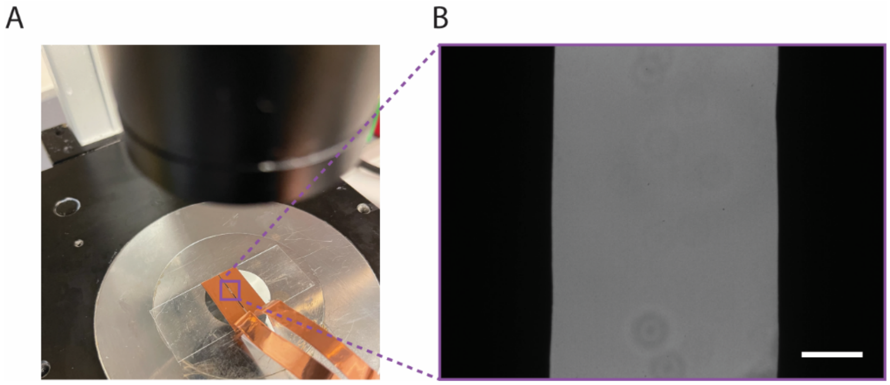


**Fig. S1.** Electrodeformation chamber. (A) Electrodefomation chamber made using copper tapes that are parallelly spaced and uniformly adhered to a coverslip glass. (B) Electrodeformation chamber image acquired using a 20X objective. Scale bar is 50 μm.

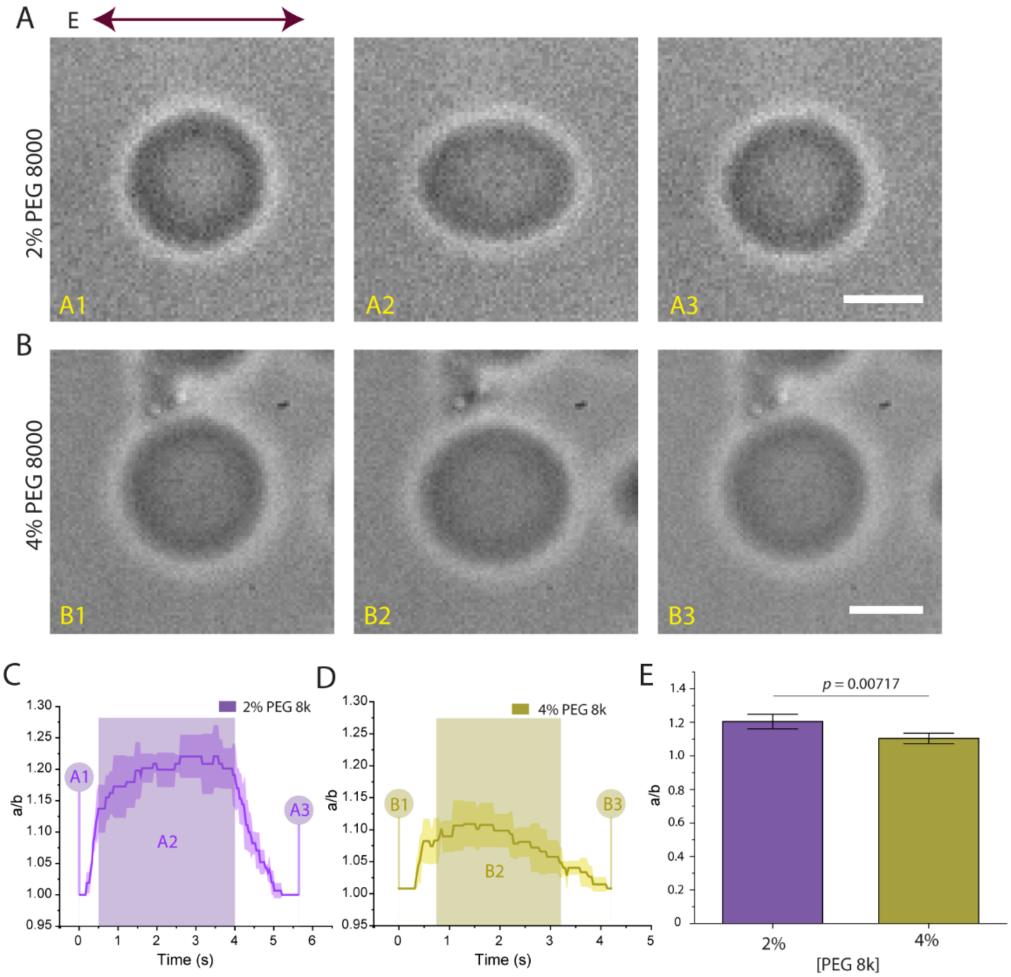


**Fig. S2.** Electroperturbation of GUVs at variable viscosity contrast ($\eta$) with a fixed conductivity ratio $\Lambda$. (A) A sequence of bright field images shows transformation of 2% PEG 8000 encapsulating GUVs from spherical (A1) to prolate deformed (A2) back to spherical recovery (A3). (B) Electrodeformation of 4% PEG 8000 encapsulating GUVs. Conductivity ratio $\Lambda$ was matched to that of 2% PEG 8000 by addition of 75 mM NaCl. (C,D) Deformation profile of 2 or 4% PEG 8000-containing GUVs in response to 30 kV/m AC field. (E) Comparison and statistical analysis of maximum GUV deformation of each GUV condition as indicated. Data represent mean maximum deformation and error bars denote ± SD. $N_{2\%}$= 10, $N_{4\%}$ = 10. (Scale bars, 10 μm)

**Table S1.** Appendix of parameters used in numerical analysis of vesicle electroperturbation

| Parameters | Description |
|---|---|
| $\varepsilon_{ex}$ | Inner solution permittivity |
| $\varepsilon_{in}$ | Outer solution permittivity |
| $\sigma_{ex}$ | Outer solution conductivity |
| $\sigma_{in}$ | Inner solution conductivity |
| $\mu_{in}$ | Inner solution dynamic viscosity |
| $\mu_{ex}$ | Outer solution dynamic viscosity |
| $C_m$ | Membrane capacitance |
| $a$ | Vesicle radius |
| $E_o$ | Electric field strength |
| $\omega$ | Frequency |
| $G_m$ | Membrane conductivity |
| $\kappa$ | Membrane bending modulus |

**Table S2.** List of dimensionless parameters

| **Dimensionless parameter** | **Equation** | **Description** | **Value for (PEG8000 2%, 4%, 8%)** |
|---|---|---|---|
| β | $\varepsilon_{ex}E_o^2 aC_m/\mu_{ex}\sigma_{ex}$ | Electric field strength | 0.99162 |
| χ | $C_m\kappa/\sigma_{ex}\mu_{ex}a^2$ | Bending rigidity | 0.559x$10^{-4}$ |
| G | $aG_m/\sigma_{ex}$ | Membrane conductivity | 0 |
| α | $\varepsilon_{ex}/aC_m$ | Bulk charge relaxation time | 7.1x$10^{-3}$ |
| Λ | $\sigma_{in}/\sigma_{ex}$ | Conductivity ratio | 0.93, 0.79, 0.65 |
| η | $\mu_{in}/\mu_{ex}$ | Viscosity ratio | 1.05, 3.02, 6.94 |
| ξ | $\varepsilon_{in}/\varepsilon_{ex}$ | Dielectric permittivity ratio | 1.01 |
| Ω | $\omega aC_m/\sigma_{ex}$ | AC field frequency | 2.79 |

**Supplementary Movie**

**Movie S1 (separate file).** Bright-field image series of electrically perturbed GUVs acquired using a high-speed camera at 400 fps. Prolate deformation of GUVs achieved by encapsulating solution with electrical conductivity ratio $\Lambda > 1$ and applying 30 kV/m AC field at 5 kHz frequency. Scale bar, 10 μm.

**Movie S2 (separate file).** Bright-field image series of oblate GUV deformation in response to electrical perturbation. Oblate deformation mode was achieved by encapsulating solution with electrical conductivity ratio $\Lambda < 1$ and applying 30 kV/m AC field at 50 kHz frequency. Scale bar, 10 μm.

**Movie S3 (separate file).** Confocal image series of electrically perturbed actin filament GUVs acquired every 170 ms. Green, ATTO 488 actin. Scale bar, 10 μm.

**Movie S4 (separate file).** Bright-field image series of 2% PEG 8000 encapsulating GUVs acquired using a high-speed camera at 400 fps. Scale bar, 10 μm.

**Movie S5 (separate file).** Electrodeformation of GUVs encapsulating 4% PEG 8000. Scale bar, 10 μm.

**Movie S6 (separate file).** Electrodeformation of GUVs encapsulating 8% PEG 8000. Scale bar, 10 μm.
